# LINGO3 interacts with Trefoil factor 2 to enforce mucosal barrier integrity and drive tissue repair during colitis

**DOI:** 10.1101/469684

**Authors:** Kelly M. Zullo, Yingbiao Ji, Yun Wei, Karl Herbine, Nicole Maloney, Rachel Cohen, Christopher Pastore, Ma Samsouk, Sriram Srivatsa, Li Yin Hung, Michael H. Kohanski, Noam A. Cohen, De’Broski R. Herbert

**Affiliations:** Department of Pathobiology, University of Pennsylvania School of Veterinary Medicine, Philadelphia, PA 19104; Department of Pathology, University of California, San Francisco, San Francisco, CA 94143, USA.; Department of Otorhinolaryngology—Head and Neck Surgery, Perelman School of Medicine at The University of Pennsylvania, Philadelphia, PA 19104; The Corporal Michael J. Crescenz VA Medical Center Surgical Service, Philadelphia, PA 19104; Monell Chemical Senses Center, Philadelphia, PA 19104; Department of Medicine, Division of Experimental Medicine, University of California, San Francisco, San Francisco, CA 94143, USA.

**Keywords:** intestinal epithelial cells, colitis, intestinal stem cells

## Abstract

Mucosal epithelia are constantly exposed to damaging stimuli from mechanical, chemical, or biologic entities, and depend on rapid repair mechanisms to maintain tissue homeostasis and immunological quiescence. The reparative cytokine Trefoil factor 2 (TFF2) serves to enforce mucosal barrier integrity, but whether TFF2 receptor(s) exist is controversial. Herein, we demonstrate leucine rich repeat and immunoglobulin like domain containing nogo receptor interacting protein 3 (LINGO3) is a necessary transmembrane component for TFF2-mediated ERK signaling, proliferation, and recovery of trans-epithelial resistance of primary epithelia during wound healing. Human respiratory and intestinal epithelia express LINGO3 and mice lacking *Lingo3* have impaired intestinal barrier function and fail to recover from DSS-induced colitis. Compared to wild-type controls, LINGO3 deficiency impairs both crypt regeneration and expression of the intestinal stem cell marker *Lgr5*. Importantly, *Lingo3*^-/-^ mice display a phenotype similar to that previously reported for *Tff2* deficiency, with increased paracellular permeability, and significant accumulation of mucosal CD4^+^ T_H_1 cells expressing IFNγ^+^ TNF^+^ even under steady-state conditions. Combined, these data reveal a previously unrecognized role for LINGO3 as a putative TFF2 receptor that regulates mucosal barrier integrity and GI inflammation.

**Significance:** Intestinal epithelial cells (IEC) are necessary for maintenance of homeostasis, resistance to infectious organisms, and overall organismal health. When injured the IEC and the underlying stroma and progenitor cell pool undergo restitutive and regenerative processes driven by the reparative cytokine Trefoil factor 2 (TFF2). This report identifies a novel receptor component for TFF2 signaling expressed on both human and mouse epithelial cells that is necessary for barrier integrity at the steady state and during colitic disease. The discovery of a novel TFF2-LINGO3 axis sheds new light on the processes controlling tissue repair, restitution, and regeneration at the mucosal interface.

## Introduction

Diverse epithelial cell lineages form a dynamic protective barrier to the external environment and form a semi-permeable shield for the body against mechanical, chemical, and organismal stimuli. Tissue injury, particularly in the gastrointestinal (GI) tract initiates mobilization of epithelial progenitors in the crypt stem cell niche, and changes in the gene expression profiles of enterocytes, M cells, goblet cells, enteroendocrine cells, Paneth cells, and tuft cells (1-3). Epithelial secretions ranging from cytokines to reactive nitrogen and oxygen intermediates and lipid mediators protect mucosal tissues against invasion by organisms to curtail inflammation and promote repair, but the knowledge of these soluble mediators is incomplete (3-5). Notably, the Trefoil factor family (TFF1, TFF2, TFF3) peptides drive largely enigmatic pathways of tissue repair and immunoregulation at the mucosal interface (6). Although TFFs are constitutively present within mucus secretions and regulate epithelial biology through restitution, cellular migration, regulation of tight junctions, and inhibition of epithelial cell apoptosis, how these molecules exert these functions is largely unknown (7-10).

Whether TFFs (TFF1-3) require receptor-mediated signal transduction events or act indirectly through modifying the rheological properties of mucus is an unsolved matter (6, 11, 12). In support of the receptor hypothesis, CXCR4, (the stromal derived factor 1 (SDF-1) receptor), also responds to TFF2 at high concentrations (13). Over-expression of TFF2 competed with SDF1α for calcium mobilization in Jurkat T cells and TFF2 required CXCR4 to phosphorylate AKT (13). Curiously however, these data do not show any interactions between TFF2 and CXCR4, and, to date, no data from *in vivo* experimentation indicates that CXCR4 deficiency recapitulates TFF2 deficiency. We and others have shown that TFF2 deficient mice have worsened disease post dextran sodium sulfate (DSS) colitis, increased pro-inflammatory cytokine secretion IFNγ, IL-6, IL-1β, and inducible nitric oxide synthase (iNOS) as well as defective mucosal barrier integrity under baseline conditions, but no such data have been reported for CXCR4 deficiency (14-16). Thus, we sought to identify and characterize a novel extracellular transmembrane protein that would be responsible for TFF2 activity and where loss of function would recapitulate TFF2 deficiency *in vivo*.

This study provides supportive evidence that LINGO3 is at least one component of a TFF2 receptor complex. Our conclusion is based on results from immunoprecipitation, confocal microscopy, and gene-deletion studies in both transformed and primary epithelial cultures. Human sinonasal and rectosigmoid tissue stained intensely for LINGO3 in the vicinity of TFF2-producing goblet cells. CRISPR-mediated *Lingo3* gene deletion in an intestinal epithelial cell line (MC38) impaired TFF2-induced ERK phosphorylation and proliferation and *Lingo3* deficient mice were phenotypically similar to reported defects in *TFF2* deficient animals in regards to mucosal accumulation of pro-inflammatory cytokines and impairment of crypt regeneration following DSS-induced colitis. Combined, these data reveal that LINGO3 is a previously unrecognized regulator of mucosal epithelial cells in humans and mice that may function as a receptor component for TFF2 signaling.

## Results

### Identification of LINGO3 as a mucosal epithelial protein in mice and humans

Despite reports that the CXCR4 antagonist AMD3100 can block rTFF2-mediated signal transduction in transformed epithelial and lymphocyte cell lines (13) there have been no reports validating a TFF2-CXCR4 axis *in vivo*. Therefore, we employed the TRICEPs approach (17, 18) using human rTFF2 as bait in order to identify potentially novel extracellular membrane-binding proteins. Preliminary results from this screen led us to focus on the orphan receptor leucine rich repeat and immunoglobulin like domain containing nogo receptor interacting protein 3 (LINGO3) as a potential TFF2 binding protein (Fig. 1A). To study LINGO3 in greater detail, a CRISPR-CAS9 gene editing approach was used to generate gene deficient mice. *Lingo3*-specific sgRNAs were generated commercially to target the translational start site in exon 2 and injected into C57BL/6 blastocysts with CAS9 protein. A founder line with a heterozygous 70 bp deletion downstream of initiator ATG was identified, bred to homozygosity. Naïve wildtype (WT) and *Lingo3* knock out (*Lingo3*KO) mice were used for Taqman assays to determine the Lingo3 mRNA expression pattern. Comparison of the intestinal epithelial cells (IEC) between WT and *Lingo3*KO mice revealed a strong signal in WT IECs that was abrogated in the *Lingo3*KO IECs, while, no clear expression was detected in splenic or lymph node populations of magnetic bead sorted CD4^+^ (T cells), B220^+^ (B cells), or CD11b^+^CD11c^+^ (myeloid cells) (Fig. 1B). *Lingo3* mRNA transcripts were detected at high levels in pulmonary and GI tract from WT mice compared to *Lingo3*KO mice (Fig. 1C). To determine whether LINGO3 had an expression pattern consistent with interaction with goblet cell-derived TFF2, combined immunofluorescence staining approach was taken using anti-human LINGO3 monoclonal antibody (mAb) and anti-human TFF2 polyclonal antibody (pAb) using formalin fixed paraffin embedded (FFPE) clinical specimens from normal subjects. In rectosigmoid tissues, TFF2 (red) localized to goblet cells located within intestinal crypts, whereas LINGO3 (green) was expressed broadly throughout colonic enterocytes and sparsely within cells at the crypt base with no staining in the isotype-matched Ab controls (Fig. 1D-I). To evaluate respiratory tissue expression, nasal polyp tissue was obtained from patients with chronic rhinosinusitis undergoing surgical resection and immunostained revealing widespread LINGO3 (green) expression on both the apical and basolateral surface of epithelia that was spatially distinct from the punctate TFF2 (red) staining inside cells with goblet cell morphology, with no staining in the isotype matched Ab control (Fig. 1J-N) Taken together, these data revealed that in human and mouse tissue, LINGO3 is predominately expressed on mucosal epithelia in the vicinity of TFF2 secretion.

**Fig. 1.**
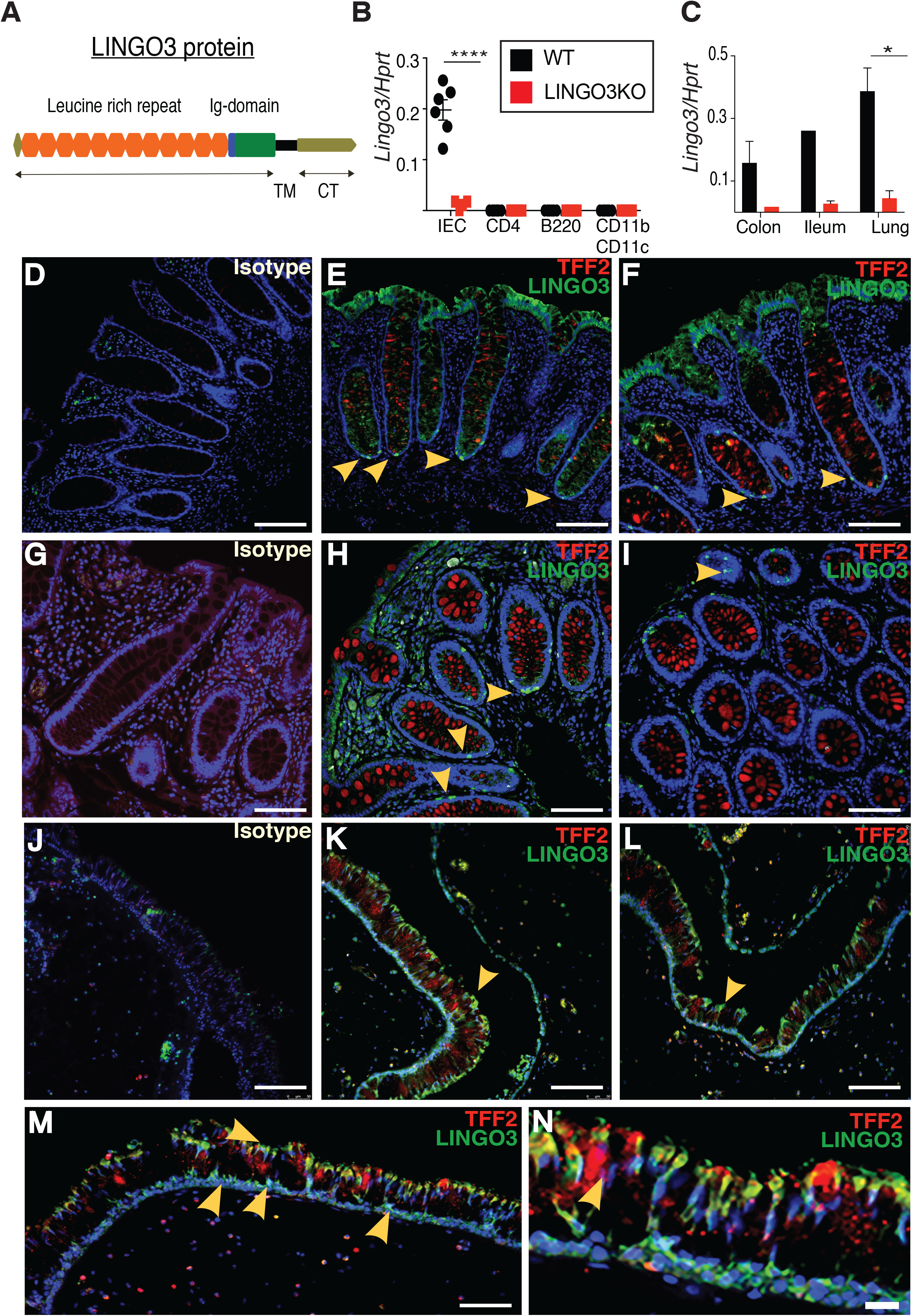
LINGO3 is a novel leucine rich repeat receptor expressed on mucosal epithelia of mice and humans. (A) Cartoon image of LINGO3 protein (B) *Lingo3* expression in intestinal epithelial cells (IECs), CD4^+^, B220^+^, CD11b^+^ CD11c^+^ cells (C) *Lingo3* expression in mucosal tissues (D-I) TFF2 (red) and LINGO3(green) immunofluorescence of human rectum (J-N) TFF2 (red) and LINGO3(green) immunofluorescence of human nasal polyps. Means ± SEM from n=4-6 samples shown. 3 independent experiments performed. *= <0.05. ****=<0.0001

### TFF2 functionally interacts with LINGO3 to promote epithelial proliferation and repair following injury

If TFF2 was a ligand for LINGO3, then several key features should be evident such as: 1) localization on the cell membrane, 2) MAPK signal transduction, and 3) induction of TFF2 biological responsiveness such as proliferation and/or resistance to apoptosis (13). To determine whether LINGO3 would localize with TFF2 at the cell membrane, RFP was cloned into the c-terminus of mouse TFF2 cDNA and transfected into human embryonic kidney 293 cells (HEK293) along with GFP-tagged mLINGO3. At 48 hr post-transfection, single or double transfected cells were identified by confocal microscopy. Whereas TFF2 single transfectants showed a punctate intra-cellular pattern (Fig. 2A), LINGO3 single transfectants localized to the cell membrane (Fig. 2B). In contrast, TFF2-RFP/LINGO3-GFP double-transfected cells, revealed a yellow signal, suggestive of co-localization in TFF2/LINGO3 conditions (Fig. 2C-D). To determine whether this co-localization also occurred with CXCR4, GFP-tagged CXCR4 was co-transfected with TFF2-RFP. Data show that TFF2-RFP remained sparse and punctate, whereas CXCR4-GFP localized to the cell surface (Fig. 2E and Fig. S1). Quantification of TFF2-RFP, Lingo3-GFP, and TFF2-RFP/LINGO3-GFP revealed that of transfected cells that were fluorescent, approximately 63% were co-transfected and of these cells all showed colocalization between TFF2-RFP and LINGO3-GFP (Fig. 2F). As an alternative approach, Chinese hamster ovary cells (CHO) were transfected with Flag-tagged cDNA encoding murine *Lingo3* for 72 hours (hrs), lysed and incubated with either TFF2-Fc or Fc, followed by Protein G sepharose immunoprecipitation and Flag-specific immunoblotting. Cell lysates from Flag-LINGO3 transfected CHO cells were incubated with either TFF2-Fc or empty vector-Fc revealing that TFF2-Fc, but not Fc, co-precipitated Flag-LINGO3 (Fig. 2G), which collectively indicated that TFF2 bound LINGO3.

**Fig.2.**
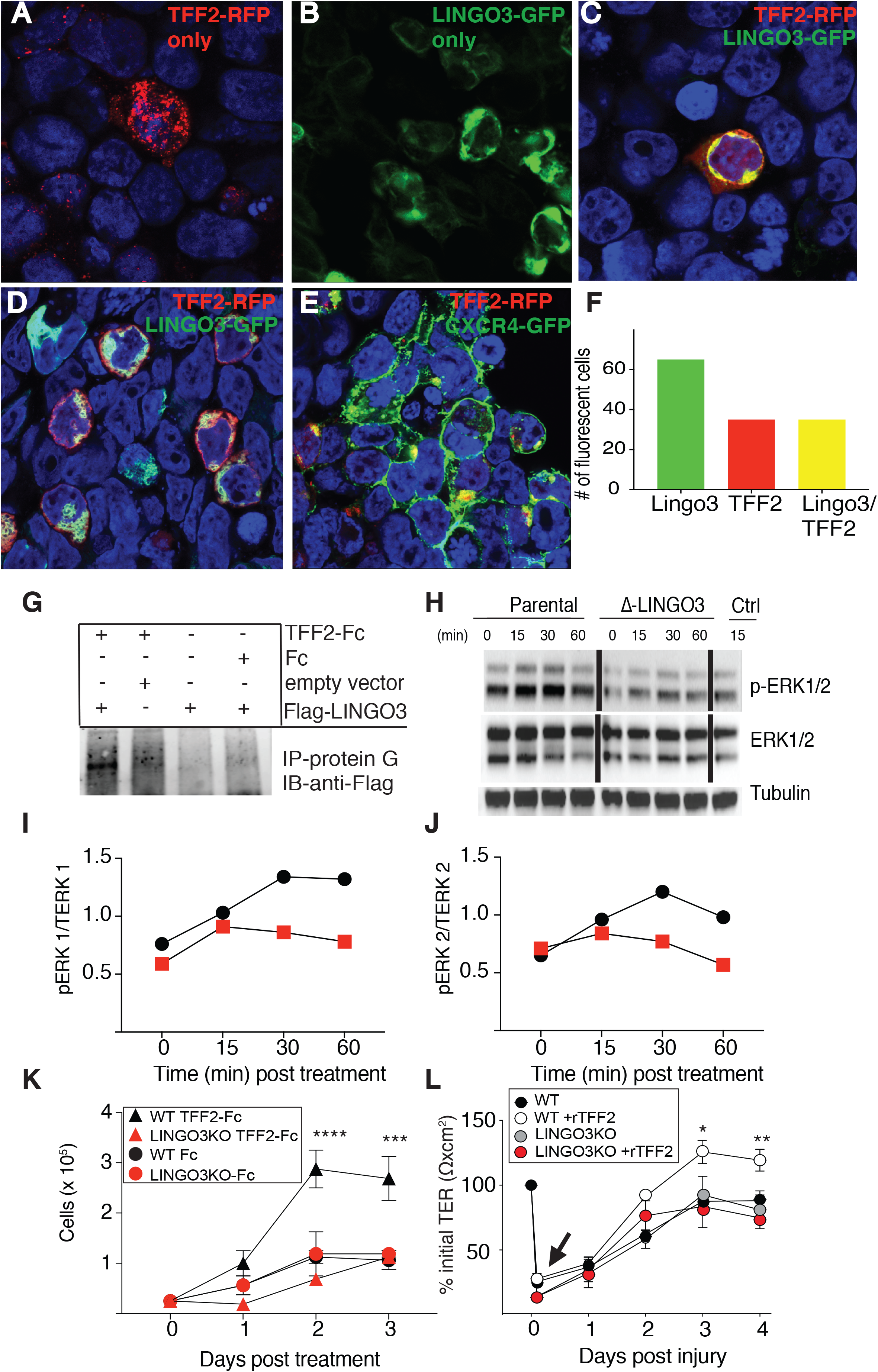
TFF2 binds and mediates downstream signaling through LINGO3. (A) Confocal images of TFF2-RFP transfected HEK293 cells (B) Confocal images of Lingo3-GFP transfected HEK293 cells (C) Colocalization of TFF2-RFP and Lingo3-GFP HEK293 transfected cells (D) Colocalization of TFF2-RFP and Lingo3-GFP HEK293 transfected cells (E) Colocalization of TFF2-RFP and CXCR4-GFP HEK293 transfected cells (F) Quantification of Lingo3, TFF2, or Lingo3/TFF2 transfected cells (G) Coimmunoprecipitation Flag-Lingo3 +/- TFF2-Fc or Fc only in CHO cells. (H) Phospho-Erk (pERK) 1/2 total ERK (ERK) 1/2, Tubulin western blot post 15, 30, 60 mins of TFF2-FC or FC only. (I) Densitometry pERK1/Total ERK1 (TERK1) J) Densitometry pERK2/TERK2 (K) Cell concentration (per mL) over three days following TFF2-Fc or Fc treatment (L) Transepithelial resistance (TEER) of primary WT and *Lingo3KO* nasal epithelia following rTFF2 (10μg/ml) or PBS treatment. Mean ± SEM from n=4-6 samples shown. 3 independent experiments. * p ≤ 0.05, ** p ≤ 0.005, *** p ≤ 0.0005, **** p ≤ 0.0001.

Next, given that TFF2 is known to induce phosphorylation of MAPK family members (19), we tested whether LINGO3 was responsible for TFF2-mediated ERK1/2 signal transduction using the MC38 murine intestinal epithelial cell line. Due to LINGO3 expression in these cells, the parental line was gene edited using CRISPRCAS9 mediated insertion of a H2-K^K^ (MHC allele) selection cassette followed by fluorescence activated cell sorting (FACS) purification and quantitative real time PCR (qRT-PCR) to confirm gene disruption (Δ-*Lingo3*) (Fig. S2). Whereas the parental MC38 line exposed to TFF2-Fc (1μg/ml) underwent increased pERK1/2 signaling with a peak at 30 min, the TFF2-Fc treated Δ-*Lingo3* MC38 line showed a moderate increase in pERK1/2 levels over control treatment with Fc protein at the time-points evaluated (Fig. 2H). Densitometric quantification confirmed the parental MC38 line had 2-fold higher p-ERK1/2 levels in response to TFF2 than Δ-*Lingo3* MC38 (Fig. 2I-J). Consistent with this signaling defect, we asked whether the loss of LINGO3 affected the ability of TFF2 to drive proliferation. Exposure of parental MC38 to TFF2-Fc (1μg/ml) under limiting serum conditions (2.5%) led to a marked increase in the cell number at 48 and 72 hrs after plating compared to Δ-*Lingo3* MC38. Further, Fc treatment did not lead to any difference in cell number of either parental or Δ-*Lingo3* MC38 cells (Fig. 2K).

If LINGO3 was responsible for TFF2 responsiveness, epithelial cells lacking LINGO3 should be deficient in restitution. Primary sinonasal epithelial cells were grown under air-liquid interface conditions in 24 transwell plates (0.4μM pore, 6.5mm diameter) (20) and subjected to scratch wound assays (21) in the presence or absence of exogenous recombinant TFF2 (rTFF2). Trans epithelial electrical resistance (TEER) was used as a surrogate for epithelial barrier integrity of primary sinonasal epithelia grown from WT vs. *Lingo3*KO mice. Following scratch with a sterile pipet, whereas the rate of TEER rebound in WT cultures was significantly augmented by the presence of rTFF2 at 3- and 4-days post-injury, there was no impact of rTFF2 treatment on *Lingo3*KO epithelia at these time-points (Fig. 2L). Thus, in both transformed and primary epithelial cultures, LINGO3 was required for the full biological effects of TFF2.

### Lingo3 deficiency increases T_H_1 cell frequency and disrupts mucosal barrier integrity

It is widely accepted that mucosal epithelia and microbial communities are in a bi-directional communicative system, with perturbations in either component resulting in inflammation-associated tissue pathology (22, 23). Because lack of TFF2 responsiveness in the absence of LINGO3 could potentially disrupt intestinal barrier integrity, cohorts of WT and *Lingo3*KO mice were inoculated orally with 440mg/kg of FITC-dextran (4 kDa) and bled 4 hrs later to measure serum levels of fluorescence as a surrogate measure of intestinal permeability. FITC-dextran levels were 3-fold increased in *Lingo3*KO mice compared to WT controls (Fig. 3A). To evaluate whether enhanced intestinal permeability was associated with alterations to the intestinal microbiome, fecal pellets were collected from naïve WT and *Lingo3*KO mice housed separately and evaluated for fecal microbial composition using 16S rRNA gene sequencing. Data show only a moderate change in microbial composition between strains (Fig. S3). Consistent with these moderate effects in microbial composition, histological evaluation of the colon tissues between strains, did not show evidence of overt pathophysiological changes between strains (Fig. 3B).

**Fig.3.**
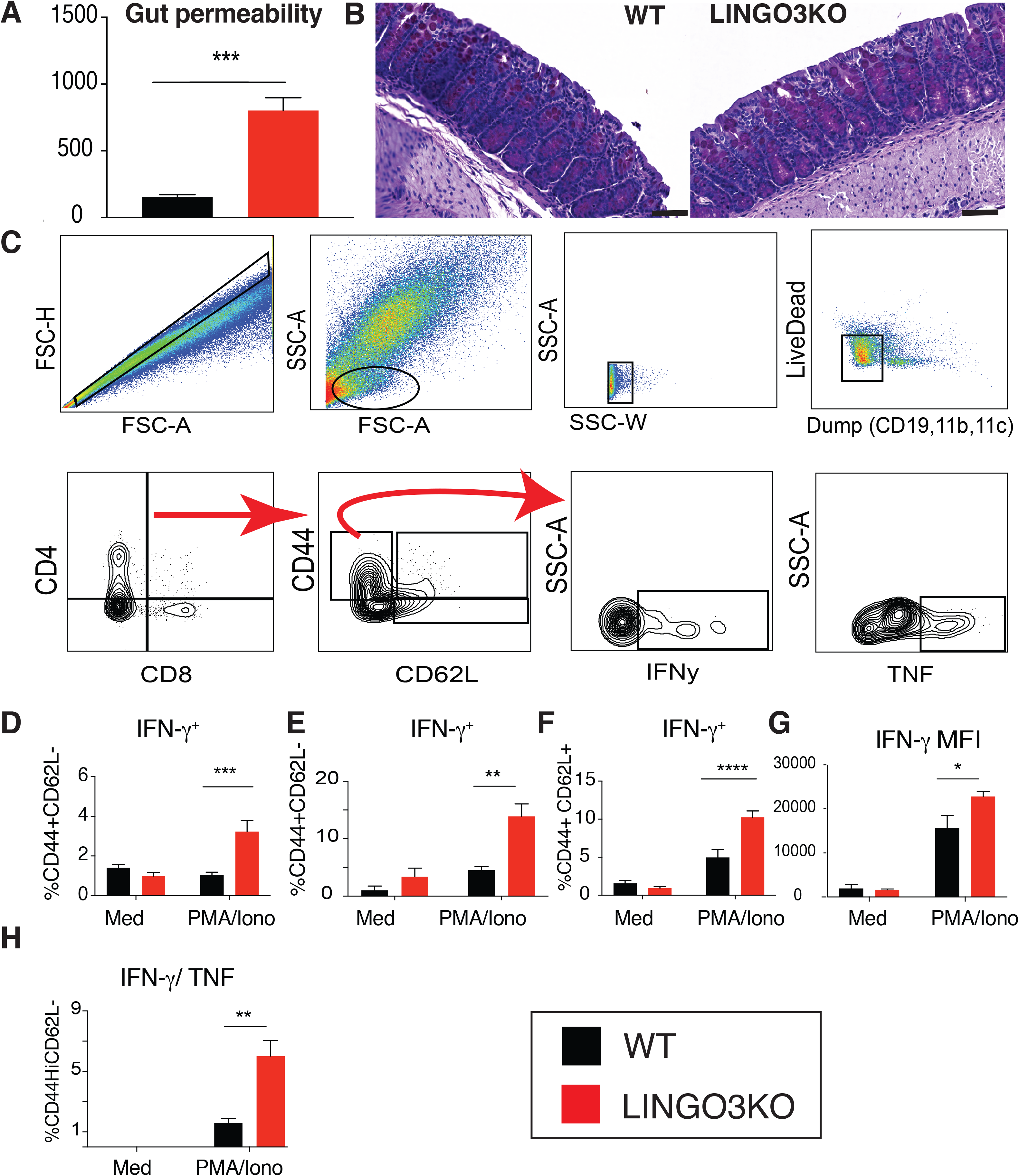
Lingo3 deficiency results in T helper 1 immune responses and intestinal permeability at the steady-state. (A) FITC-Dextran concentration (μg/mL) in serum. (B) PAS colon images from WT and *Lingo3KO* mice 17x magnification (C) Flow cytometry gating strategy for IFNγ^+^ TNF^+^ CD4^+^ T cells (D) IFNγ effector CD4^+^ cells frequencies from colon (E) IFNγ effector CD4^+^ frequencies from MLN (F) IFNγ central memory CD4^+^ cell frequencies from MLN (G) IFNγ MFI in central memory CD4^+^ cells from MLN (H) IFNγ^Hi^TNF^HI^ effector CD4^+^ frequencies in colon. Mean ± SEM from n=4-6 mice/genotype. 3 independent experiments. * p ≤ 0.05, ** p ≤ 0.005, *** p ≤ 0.0005, **** p ≤ 0.0001

However, there was a marked difference between strains in WT vs. *Lingo3*KO CD4^+^ T cells following evaluation for IFN-γ^+^ and TNF^+^ frequencies in effector (CD44^HI^ CD62L^LO^) and central memory (CD44^Hi^ CD62^HI^) populations (Gating strategy Fig. 3C). While there were no differences observed in the naïve pool (data not shown), the T cell effector pool in colon lamina propria and mesenteric lymph node (MLN) were significantly increased in *Lingo3*KO mice compared to WT mice (Fig. 3D-E). MLN CD4^+^ T cells isolated from *Lingo3*KO mice had a 2-fold increase in IFNγ^+^ memory compartment as compared to the WT controls (Fig. 3F-G). Also, IFNγ^+^TNF^+^ double positive CD4^+^T cells were significantly increased in colon tissue of *Lingo3*KO as compared to WT (Fig. 3H). Consistent with the IFNγ^+^TNF^+^ colonic CD4^+^ T cells being memory CD4^+^ T cells, 4 weeks of broad-spectrum antibiotic treatment did not reduce their frequencies despite a reduction in the microbial load by several orders of magnitude (data not shown). Furthermore, no differences were observed in effector or central memory CD8^+^ T cells (data not shown). Taken together this indicates that *Lingo3* deficiency increased the basal Type 1 cytokine response in the intestine commensurate with reduced barrier integrity.

### *Lingo3*KO mice have enhanced sensitivity to DSS induced colitis and epithelial intrinsic defect in intestinal stem cell gene expression

*Tff2* deficient mice have enhanced gut permeability and sensitivity to DSS-induced colitic injury (14, 16). We postulated that if a TFF2-LINGO3 interaction existed, *Lingo3* deficiency would recapitulate these features in the DSS model. Age matched and co-housed female WT and *Lingo3*KO mice were administered 2.5% DSS ad libitum in the drinking water over 6 days, then returned to normal drinking water and monitored for an additional 4 days for changes in weight and disease activity index (DAI). Comparison of weight change and disease activity index (i.e. hunched posture, fecal blood, and diarrhea) revealed that *Lingo3*KO mice failed to regain body weight following the cessation of DSS treatment (Fig. 4A) and also continued to show increased DAI scores compared to WT mice (Fig. 4B). Colon lengths on day 10 were shorter in *Lingo3*KO mice compared to WT (Fig. 4C).

**Fig.4.**
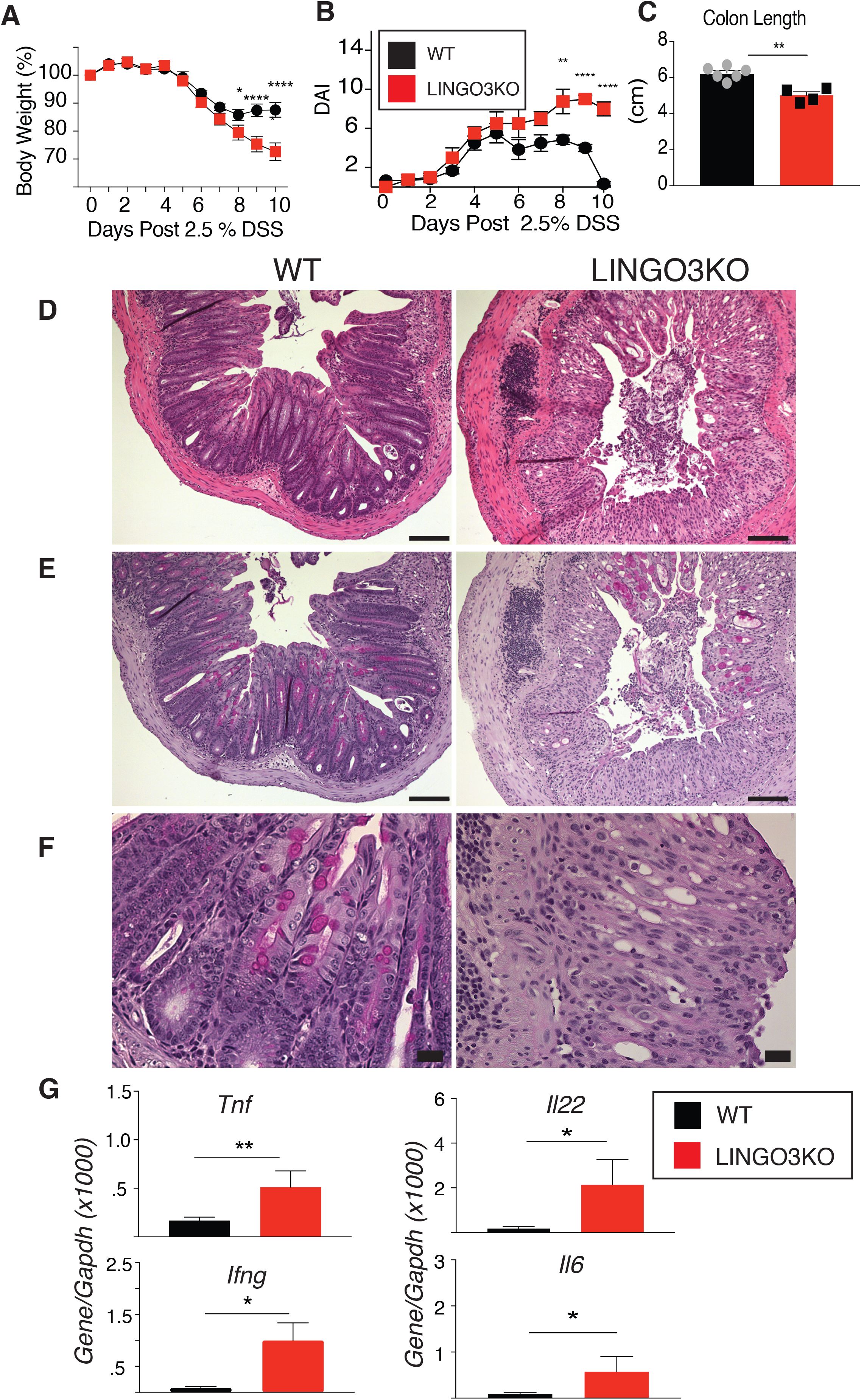
DSS induced colitis is more severe in *Lingo3KO* mice compared to WT. (A) Body weight percentage (%) during 2.5% DSS induced colitis (B) Disease activity score (DAI) during 2.5% DSS induced colitis (C) Colon length (cm) 10 days post 2.5% DSS (D) H&E images of colon day 10 post 2.5% DSS 10x (E) PAS images of colon day 10 post 2.5 DSS 10x (F) PAS images of colon day 10 post 2.5% DSS 40x (G) Pro-inflammatory mRNA expression in colon 10 days post 2.5% DSS administration. Mean ± SEM from n=4-6 mice/genotype. 3 independent experiments. *p ≤ 0.05, ** p ≤ 0.005, **** p ≤ 0.0001

Histological evaluation at day 10 revealed that *Lingo3* deficient mice had increased inflammatory cell infiltration, ulceration, submucosal edema, and smooth muscle hyperproliferation as compared to WT at day 10 (Fig. 4D). Using Periodic acid Schiff (PAS) staining to reveal mucins, we observed that PAS^+^ goblet cells rapidly re-populated the intestines of WT mice, whereas goblets cells were sparse in the intestines of *Lingo3*KO mice at this timepoint (Fig. 4E-F). Due to these differences, experiments tested whether *Lingo3* deficiency heightened the inflammatory cytokine gene expression and/or altered intestinal stem cell gene expression. Under co-housed conditions, *Lingo3*KO expressed significantly higher *Tnf*, *Il22*, *Ifng*, and *Il6* mRNA transcript levels at day 10 compared to WT (Fig. 4G).

Next, we evaluated expression levels for leucine-rich repeat-containing G-protein coupled receptor 5 (*Lgr5*), the receptor for R-spondins, that marks intestinal stem cells located at the base of the crypt (24). Comparison of *Lgr*5 expression levels between WT and *Lingo3*KO mice under steady-state and DSS-treated conditions both revealed a significant reduction in *Lgr5* expression in *Lingo3*KO (Fig. 5A). To distinguish whether reduced *Lgr5* expression was epithelial cell-intrinsic, 3D organoid cultures were generated from the small intestine. Strikingly, primary organoid cultures from *Lingo3*KO mice at day 6 showed a marked decrease in overall growth (Fig. 5B) and *Lgr5* expression as compared to WT (Fig. 5C). Collectively, these data imply LINGO3 serves a critical role in intestinal epithelial cell repair following injury.

**Fig. 5.**
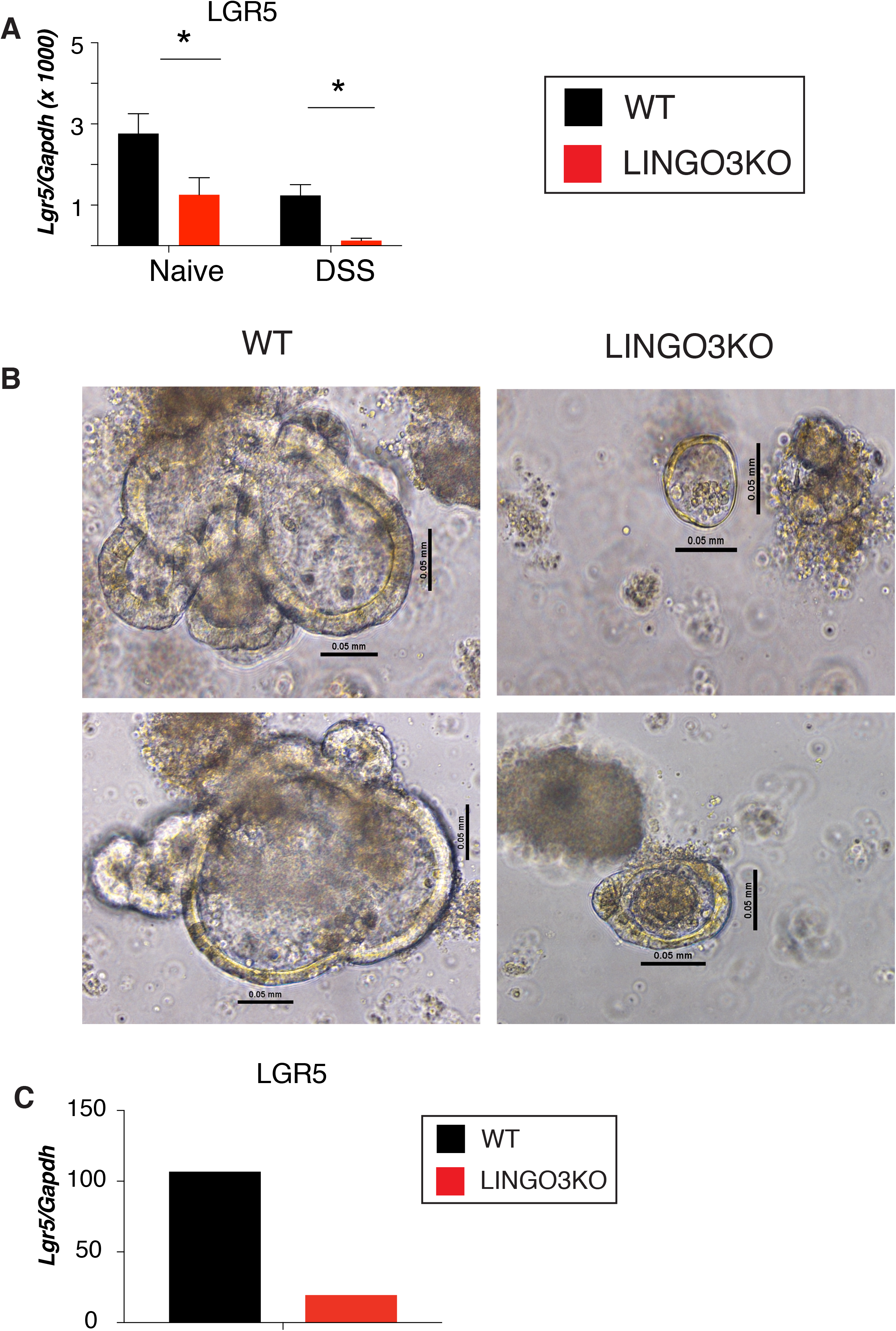
*Lingo3KO* mice have epithelial intrinsic defect in intestinal stem cell regeneration during colitis and at steady state. (A) *LGR5* mRNA expression from colons of naïve and DSS treated mice. Graphs show means ± SEM from n=4-6 mice/genotype. (B) Brightfield microscopy images day 6 of small intestinal organoid culture 20x (C) *LGR5* mRNA expression from small intestinal organoids day 7 of culture. 3 independent experiments performed. * p ≤ 0.05

## Discussion

This study provides evidence that the orphan transmembrane protein LINGO3 functions as an essential component for epithelial responsiveness to Trefoil factor 2 (TFF2). This finding is particularly important given the body of evidence indicating that TFF2 regulates epithelial repair, restitution and inflammation within the GI and respiratory tract (7, 14, 15, 25-27). LINGO3 has no known function(s) reported to date, therefore these data suggest we have uncovered a novel regulator of mucosal epithelial cell biology. In human tissues, LINGO3 was constitutively expressed on both the apical and basolateral surfaces of epithelial cells lining the sigmoid colon and turbinate sinuses of the upper airway. Loss of function studies in transformed and primary epithelial cells indicated that TFF2- driven cell expansion, recovery of barrier function after wounding and localization at the cell membrane were all impaired in the absence of *Lingo3*. Importantly, the expression of LINGO3 in the crypt intestinal stem cell (ISC) niche, combined with the marked defect in *Lgr5* pre and post DSS mediated injury indicates that LINGO3 may have important roles in intestinal regeneration.

Pursuit of novel candidate TFF receptor(s) led us to employ TRICEPs technology that was created to identify low-affinity protein interactions with glycosylated cell-surface receptors. This approach led to a focus on LINGO3. LINGO3 is a 166 amino acid (AA) single pass transmembrane protein with 11 leucine repeats, 8 N-linked glycosylation sites, a single disulfide bond, and a short cytoplasmic tail containing 2 serine’s and a single tyrosine (https://www.uniprot.org/uniprot/Q6GQU6). Previous reports have suggested that CXCR4, the canonical SDF-1 receptor, additionally serves as a low affinity receptor for TFF2 (13). To date however, there are no data showing direct binding between TFF2 and CXCR4, CXCR4 dependent signaling, or that CXCR4 deficient mice phenocopy any aspect of TFF2 deficiency such as impaired barrier integrity, basal Type 1 inflammation, or defects in the gastric compartment (16, 28). We demonstrate that TFF2 co-immunoprecipitated with LINGO3 and localized to the outer rim of LINGO3 transfected cells, whereas this co-localization pattern was not observed with CXCR4 transfectants. In addition, the TFF2 driven phosphorylation of ERK that has been demonstrated in bronchial and gastric epithelial cells (13, 19) was also shown here within intestinal epithelia. Most importantly however, restitution, which is the ordered movement of cells to repopulate denuded basement membrane, is a hallmark of TFF function. Our data show that following pipet-mediated scratch wounding, TFF2-Fc administration significantly accelerated the rebound in TEER of WT, but not *Lingo3* deficient epithelia. However, TEER eventually recovered in all groups, indicating that LINGO3 only partially controls epithelial barrier recovery. Nonetheless, *in vivo*, LINGO3 deficiency significantly increased intestinal permeability under steady-state conditions, which was strikingly consistent with *Tff2*KO mice (16).

Increased intestinal permeability is potentially life-threatening due to TLR-driven inflammatory responses. Indeed, CD4 lymphocytes producing both IFN-γ and TNF were markedly increased in naïve *Lingo3*KO compared to co-housed WT controls. These features are similar to those of *Tff2*KO mice (16) and further support our overall contention that TFF2 and LINGO3 coordinately function to control mucosal barrier integrity, which if lost, can affect subsequent responses against pathogens or tissue damaging agents. *Lingo3*KO mice were more susceptible than WT to the tissue damaging effects of DSS. Importantly, the protective role for LINGO3 was evident during the healing phase following cessation of DSS treatment. Furthermore, *Lingo*3KO mice failed to re-populate their intestinal goblet cells at the time when these cells were clearly repopulating the intestines in their WT co-housed counterparts.

Unexpectedly, we found that in human rectal tissue, LINGO3 was constitutively expressed in pattern consistent with the location of the intestinal stem cell niche (ISC) where LGR5^+^ cells are found. LINGO3 deficiency led to an intrinsic defect in *Lgr5* expression both *in vivo* and *ex vivo* within 3-D intestinal organoids, suggesting that hematopoietic cell-derived inflammatory cytokines were not the cause of reduced *Lgr5* expression seen in intact animals. Following crypt irradiation, LGR5^+^ stem cells respond to intestinal damage by repopulating the intestine (1, 24, 29). Although there have been no implications for TFF proteins in the maintenance of the intestinal stem cell niche, given the localization of LINGO3, the interaction between TFF2 and LINGO3, and the role of ISC niche in regeneration, it is possible that a TFF2/LINGO3 axis serves an important role in regulating the ISC niche to augment repair under steady-state or injurious conditions.

It is unlikely that LINGO3 acts alone to facilitate TFF2 responsiveness given evidence that the related family member LINGO1 is one subunit of a trimolecular complex including TROY and nogo receptor (NgR)(30, 31). Indeed, *Lingo3* deficient epithelial cells have impaired ERK phosphorylation, but not a total loss of ERK phosphorylation. Thus, the effect of LINGO3 deficiency on the GI tract may only partially reflect the importance of TFF biology in homeostasis and regenerative repair, particularly due to redundancies in TFF function. In sum, we propose that LINGO3 is a novel regulator of TFF2 responsiveness that mediates epithelial proliferation, signaling and barrier integrity, which may also be evident at mucosa barriers other than respiratory and intestinal tract.

## Author contributions

K.M.Z., Y.J., Y.W., K.H., N.M., R.C., C.P., M.H.K., and L.Y.H., performed experiments, K.M.Z., Y.J., M.H.K, S.S., N.A.C., and D.R.H, analyzed data, M.S., provided reagents, D.R.H. conceived the study and K.M.Z. and D.R.H, wrote the manuscript.

## Acknowledgements

D.R.H. is supported by NIH (AI095289, GM083204, UO1AI125940) and the Burroughs Wellcome Fund. NAC is supported by NIH (DC013588).

## Experimental Procedures

### Mice

All animal procedures were approved by the Institutional Animal Care and Use Committee at University of California, San Francisco. Super-ovulated female C57BL/6 mice (4 weeks old) were mated to C57BL/6 stud males. Fertilized zygotes were collected from oviducts and injected with Cas9 protein (30 ng/ul), sgRNA (15 ng/ul) into pronucleus of fertilized zygotes. Injected zygotes were implanted into oviducts of pseudopregnant CD1 female mice. Animals were housed under specific-pathogen free barriers in vivarium at San Francisco General Hospital or University of Pennsylvania. All the procedures were reviewed and approved by IACUC at University of California at San Francisco (protocol #AN109782-01) and University of Pennsylvania (protocol #805911).

### Dextran sodium sulfate (DSS) model

All experiments used six to ten-week-old wildtype (WT) or *Lingo3*KO C57BL/6 female mice bred in house and cohoused for four weeks. Colitis was induced in mice using 2.5% Dextran Sulfate Sodium Salt (w/v, 0216011080, MP-Biomedicals LLC Solon, OH) in autoclaved drinking water for 6 days. Fresh solution was provided on day 3, on day 6 mice were returned to normal water and monitored until necropsy. Body weight, appearance, fecal occult blood (Fisherbrand™ Sure-Vue™ Fecal Occult Blood Slide Test System, ThermoFisher Scientific, Inc., Waltham, MA, USA), stool consistency, and diarrhea were recorded daily from tatooed animals. Colon length was determined at necropsy. Disease activity index (DAI) a cumulative score with max of 12.

### Human tissue and immunostaining

Archived rectosigmoid biopsy samples were from the SCOPE cohort at the University of California, San Francisco (UCSF). The SCOPE cohort is an ongoing longitudinal study of over 1,500 HIV-infected and uninfected adults followed for research purposes. The UCSF Committee on Human Research reviewed and approved the SCOPE study (IRB# 10-01218), and all participants provided written informed consent. Human sinonasal polyp samples were obtained from patients undergoing sinonasal surgery for the management of their chronic rhinosinusitis with informed consent and full approval of the University of Pennsylvania Institutional Review Board (Protocol #800614). FFPE tissue sections were evaluated under Cy2, Cy3, and Cy5 filters on a Leica Inverted Microscope DMi8 S Platform and a Leica DM6000B microscope with an automated stage coupled with a Leica DFC350FX camera.

### Transfection and immunofluorescence

HEK293 cells were cultured in DMEM medium (Invitrogen) supplemented with 10% fetal bovine serum (Invitrogen) and 1% penicillin-streptomycin (Invitrogen) at 37°C with 5% CO_2_. The transfection was done by the application of X-tremeGENE HP DNA transfection reagent (Roche) according to the manufacture’s manual. 2 ml of cells (0.2 × 10^6^/ml) per well was seeded in a well of LabTek Chamber slide (Sigma) for transfection. A mixture of 1μg pcDNA3:mTFF2: RFP and 1μg pcDNA3:mLingo3: GFP plasmid were added to the cells along with XtremeGENE HP DNA transfection reagent (Roche). After three-days culture, fixation was performed with 4% paraformaldehyde in PBS for 10 min and stained with DAPI (0.5μg/ml) for 5 mins. After stained cells were mounted on the coverslip, Lingo3: GFP and TFF2: RFP were visualized using the Leica TCS-NT confocal microscope.

### Co-immunoprecipitation

2.4 × 10^6^ CHO cells were transfected with 24 μg pCMV6:Lingo3:Flag (MR215769, Origene) or pCMV6 entry plasmids. After incubation of the transfection mixture with the cells for 60 h, the cells were washed twice with PBS and lysed with 300 μL digitonin extraction buffer [1% digitonin; 150 mM NaCl; 1 mM MgCl_2_; 10 mM TrisHCl, pH 7.4, 1X Proteinase Inhibitor (Pierce), 1mM PMSF]. The protein concentration was measured using BCA assay kit (Invitrogen). Around 0.6 mg total protein was mixed with 20μg purified TFF2-Fc (Ray Biotech) or Fc (Ray Biotech) and precleared with control agarose resin for 1 hr at 4°C. The lysate samples were then precipitated with 40 μl protein A/G agarose (Invitrogen) overnight at 4°C. The immunoprecipitation (IP) complex was washed with TBS for 5 times and eluted with 4× SDS loading buffer at 95°C. The IP elutes along with whole cell lysates were immunoblotted with rabbit anti-FLAG (1:400) followed by incubation with goat anti-rabbit IgG Cy5 (1:1000).

### Generation of Lingo3 knockout cell line

Homologous recombination (HR) mediated CRISPR-CAS9 genome editing was used to generate a *Lingo3*KO colon epithelial cell line (MC38). Validated *Lingo3* guide RNA obtained commercially (PNA Bio). Briefly, MC38 cells were cultured in DMEM medium (Invitrogen) supplemented with 10% fetal bovine serum (Invitrogen), 1X non-essential amino acids (NEAA), 10mM HEPES, 1mM Sodium Pyruvate, 50ug/ml Gentamicin and 1% penicillin-streptomycin (Invitrogen) at 37°C with 5% CO_2_. Before transfection, 3 ml of cells (1 × 10^5^/ml) per well was seeded in a well of a six-well plate for overnight culture. 1μg validated Lingo3 sgRNA (PNA Bio), 6μg Lingo3 donor vector (PNA Bio) and 1μg Cas9 plasmid (PNA Bio) were cotransfected into the cells using X-tremeGENE HP DNA transfection reagent (Roche). After expanding the cells for the two generations, H2KK positive cell (*Lingo3KO*) cells were identified by flow cytometry because Lingo3 donor vector contains a H2KK gene cassette. Loss of *Lingo3* expression in H2-K^K^ positive cells was further confirmed by qRT-PCR analysis.

### Western Blotting

1 ml of M38 parental or *Lingo3KO* cell (1 × 10^5^/ml) per well was cultured in a well of a 24-well plate and cultured for 48 hrs in complete medium. After cells were washed with 1ml PBS and cultured in 1ml serum-free medium for 24 hrs for serum starvation the cells were treated with TFF2-Fc (1μg/ml) in the complete medium for 0, 5min, 15min, 30min and 1hr. After washing the cells with 1ml PBS twice at the indicated time, the total protein was extracted with the lysis buffer [1% Triton X-100, 150 mM NaCl,10 mM Tris (pH 7.4), 1 mM EDTA, protease and phosphatase inhibitor cocktail (Sigma)]. 15μg samples were loaded and separated in 4-12% PAGE gel (Invitrogen). The following primary Abs were used for immunoblotting assays: rabbit anti-phospho-p44/p42 MAPK (Erk1/2) (Cell signaling,1:2000), rabbit anti-p44/42 MAPK (ERK1/2) (Cell-Signaling 1:2000), mouse anti-α-Tubulin (Invitrogen, 1:2500).

### Cell growth analysis

WT and *Lingo3KO* MC38 cell lines were plated at a density of 0.25 × 10^5 cells/ well in duplicate in DMEM media containing 2.5% serum and TFF2-FC (1μg/ml) or FC (1μg/ml) added to each well.

### 16S rRNA gene sequencing

Genomic DNA was extracted from fecal pellets of mice using Qiagen DNeasy PowerSoil Kit. The V4 region of the 16S rRNA gene was amplified using barcoded primers for use on the Illumina platform (32, 33). Sequencing was performed using 250-base paired-end chemistry on an Illumina MiSeq instrument. QIIME v. 1.8 (34) was used to process paired-end reads and to perform downstream analyses including taxonomy assignment and heat map generation.

### Histological staining

Mice were euthanized and colons extracted. Colon tissue was either opened longitudinally, flushed with ice-cold PBS to remove feces and formalin fixed as “swiss rolls” or 1-2 cm of the distal colon was formalin fixed. Colon tissues were stained with hematoxylin and eosin and periodic acid Schiff stain (PAS) performed by the Veterinary comparative pathology core at the University of Pennsylvania. A Leica widefield microscope was used for imaging.

### Quantitative RT-PCR

For intestinal epithelial cell isolation, colons and small intestines were dissected from WT and *Lingo3KO* mice and cDNA was prepared. Gene expression was measured using BIO-RAD CFX machine and data normalized to GAPDH and presented as MEAN ± SEM from the replicates. For the Taqman hybridization assays the manufacturer’s protocol was used. Data was normalized to HPRT. All real-time PCR reactions were carried out on Bio-rad CFX96 system and data were analyzed using the ΔΔCt method as recommended in the manual.

### Flow Cytometry

MLN was dissected from WT and *Lingo3KO mice*, homogenized, and resuspended in 1mL of RPMI and processed for analysis as described in supplemental methods.

### Evaluation of intestinal permeability

WT and Lingo3^-/-^ C57BL6 non-cohoused and cohoused (4 weeks) mice were fasted for 16 hours and orally gavaged with 200 μL of 44mg/100g of 4kDA fluorescein isothiocyanate dextran (FITC-dextran) (Sigma, FD4). Four hours post administration, mice were euthanized and blood was collected and stored A standard curve was generated from 0.1- 1 mg/mL of FITC-dextran and fluorescence was measured at a wavelength of 520 nm using Synergy 2 microplate reader and the Gen 5.0 software (BioTek).

### Nasal ALI and intestinal organoid cultures

Primary murine nasal epithelial cells were grown for 7 days as submerged cultures and for 2 weeks at air liquid interface in transwells. On day 21, monolayers were wounded in a cross pattern with a sterile pipet tip (1mm width) and treated daily on the apical side with 10 ng/ml recombinant human TFF2 (Peprotech) or 1xPBS (Mock). Cultures were rinsed with 1X PBS for 5 minutes prior to TEER measurement. Serial measurements of TEER (Ohms-cm2) were taken using an EVOM (World precision instruments) and the cup/transwell insert adapter. Crypts were isolated from mouse small intestinal jejunum as previously reported (35, 36) with modifications indicated in supplement.

### Statistics

Statistical analyses were performed using GraphPad Prism version 7.0 (GraphPad). A two-tailed Student’s T test or ANOVA was used where appropriate.

---

**Fig. S1.**
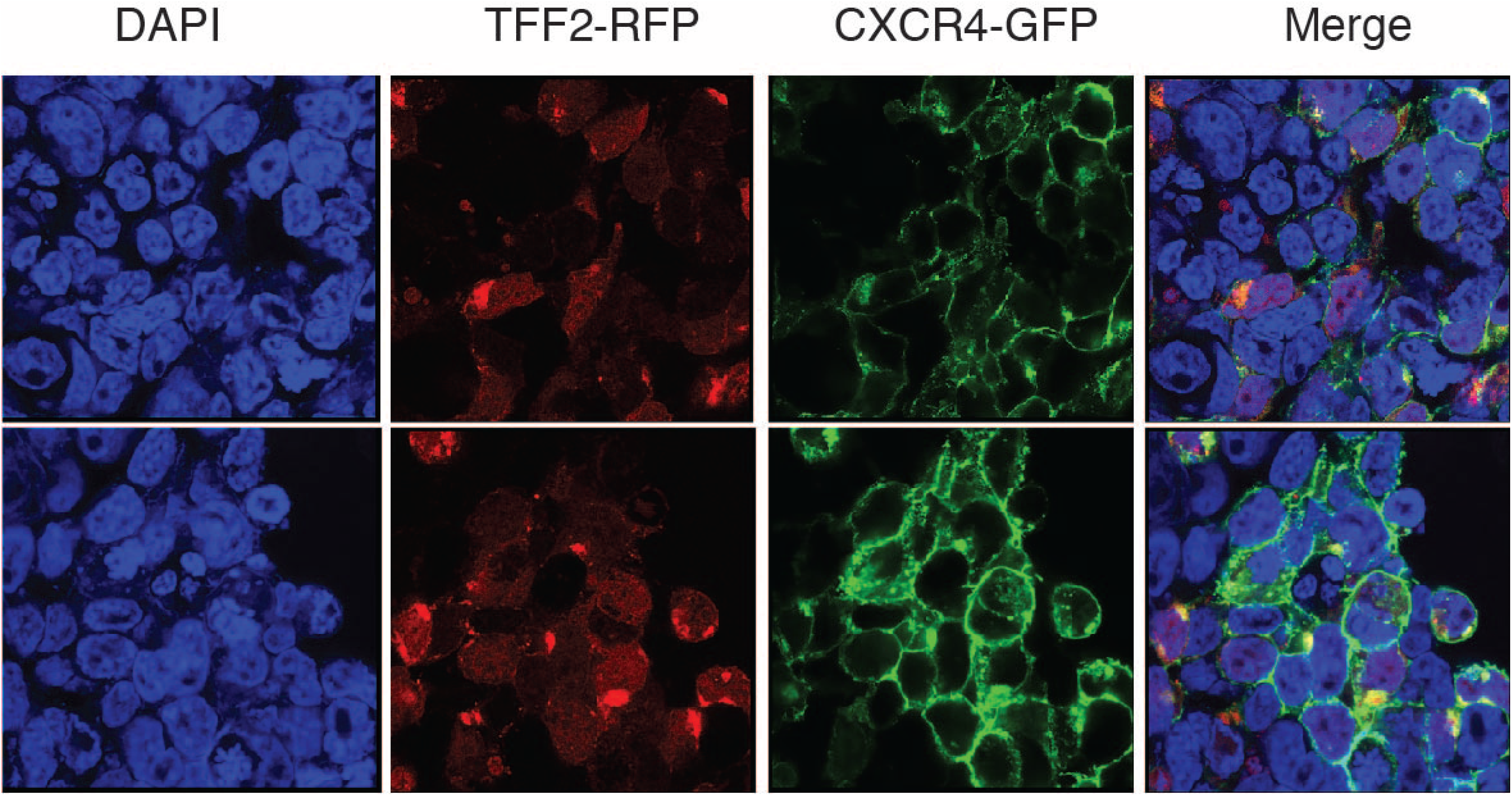
Representative single color and merged images from HEK cells transfected with TFF2-RFP and CXCR4-GFP constructs

**Fig. S2.**
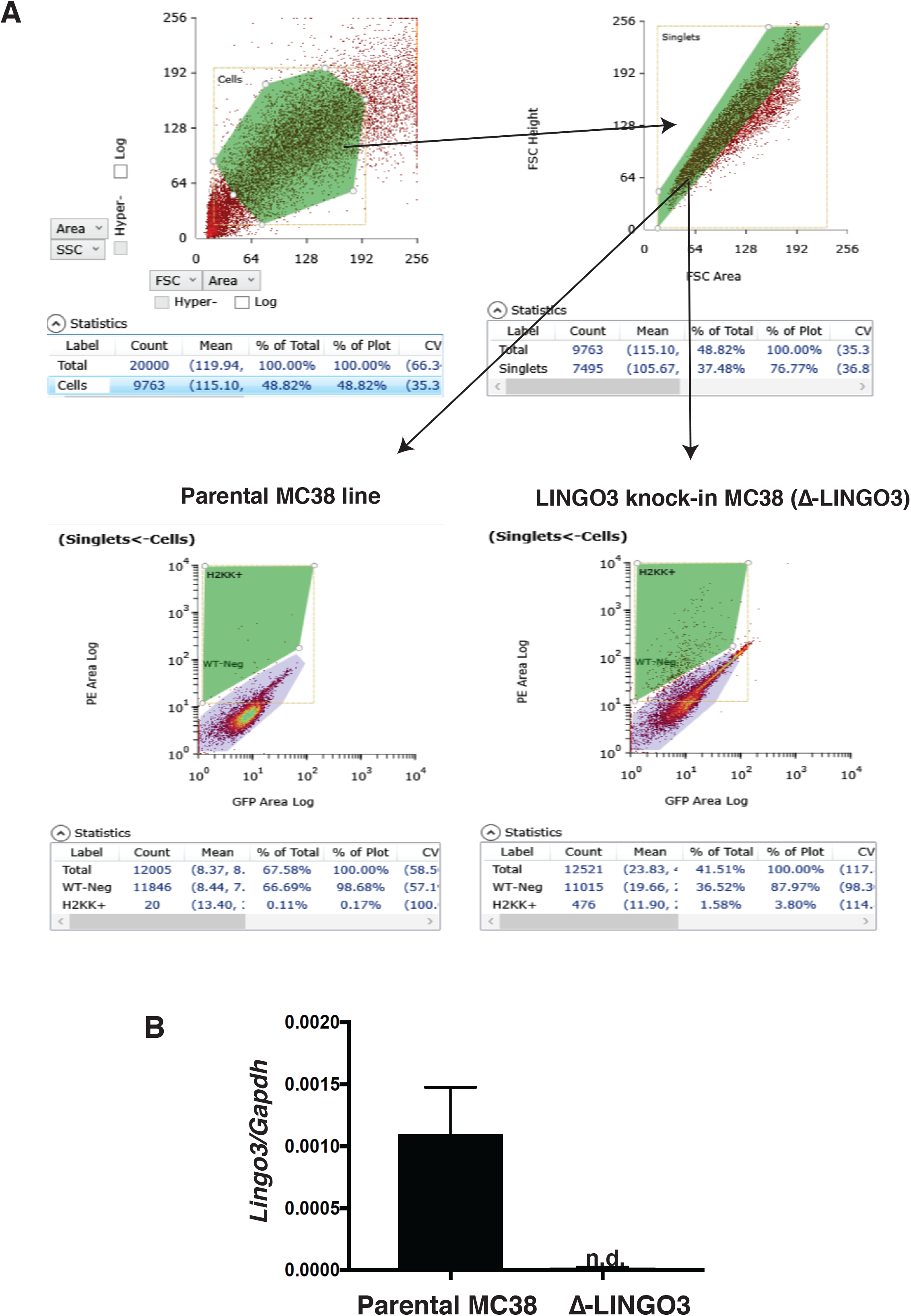
A) Flow cytometry gating strategy to identify MC38 cells that had undergone successful CRISPR/CAS9 mediated knockin of an H2K–K allele for cell sorting and subcloning. B) Comparison of Lingo3mRNA expression levels by real time PCR

**Fig. S3.**
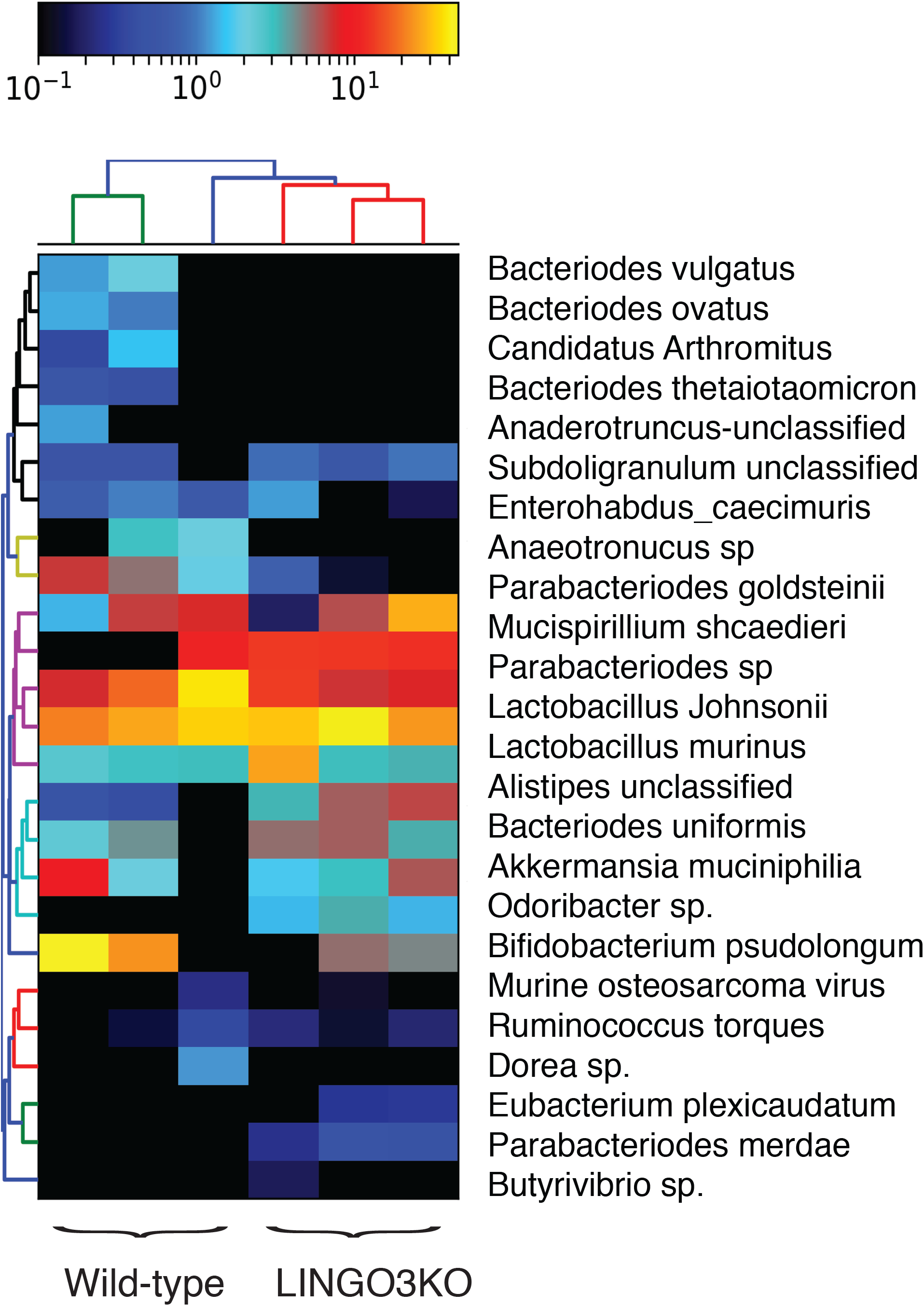
WT and LINGO3KO mice have moderate differences in microbial composition at baseline. The heat map based on 16S rRNA marker gene sequencing of fecal pellets was generated using QIIME v. 1.8. OTUs (rows) and individual mice (columns) are clustered by phylogenetic relationship.

